# DiPPI: A curated dataset for drug-like molecules in protein-protein interfaces

**DOI:** 10.1101/2023.08.09.552637

**Authors:** Fatma Cankara, Simge Senyuz, Ahenk Zeynep Sayin, Attila Gursoy, Ozlem Keskin

## Abstract

Proteins interact through their interfaces, and dysfunction of protein-protein interactions (PPIs) has been associated with various diseases. Therefore, investigating the properties of the drug-modulated PPIs and interface-targeting drugs is critical. Here, we present a curated large dataset for drug-like molecules in protein interfaces. We further present DiPPI (Drugs in Protein-Protein Interfaces), a two-module website to facilitate the search for such molecules and their properties by exploiting our dataset in drug repurposing studies. In the interface module of the website, we extracted several properties of interfaces, such as amino acid properties, hotspots, evolutionary conservation of drug-binding amino acids, and post-translational modifications of these residues. On the drug-like molecule side, we curated a list of drug-like small molecules and FDA-approved drugs from various databases and extracted those that bind to the interfaces. We further clustered the drugs based on their molecular fingerprints to confine the search for an alternative drug to a smaller space. Drug properties, including Lipinski’s rules and various molecular descriptors, are also calculated and made available on the website to guide the selection of drug molecules. Our dataset contains 534,203 interfaces for 98,632 proteins, of which 55,135 are detected to bind to a drug-like molecule. 2,214 drug-like molecules are deposited on our website, among which 335 are FDA-approved. DiPPI provides users with an easy-to-follow scheme for drug repurposing studies through its well-curated and clustered interface and drug data; and is freely available at http://interactome.ku.edu.tr:8501.

## 1. Introduction

Drug repurposing is an ingenious strategy to exploit available drugs for a use other than their original purpose (Pushpakom et al. 2019). Given the laborious and lengthy process of drug discovery studies, repurposing is promising in producing alternative solutions in a shorter time, with less cost, higher efficiency, and less toxicity (Parvathaneni et al. 2019). However, identifying new targets for repurposed drugs is not trivial (Ozdemir et al. 2019; Pushpakom et al. 2019). Drugs can bind to multiple targets with similar cavities (Duran-Frigola et al. 2017; Zhang et al. 2011; Engin et al. 2012). This fact leads to the interpretation that similar drugs may also target similar local binding sites (Adasme et al. 2021). Therefore, the search for alternative targets for a drug may be attained in similar local binding sites even when the global similarity is low. Interfaces are one such binding site, and two proteins that do not share an overall global similarity might share similar interface regions (Keskin and Nussinov 2007; Gao and Skolnick 2010). This property makes specific identification and characterization of interfaces a promising approach for finding new protein candidates, as physical binding to a target to modulate downstream mechanisms is a common way drugs exert their effects (J. Ma et al. 2019). Therefore, one of the strategies in drug repurposing is targeting protein interfaces that possess similar properties to the original target protein of a drug. Although targeting PPIs for therapeutic purposes is challenging, there has been an increasing interest in PPI drug targets thanks to the recent advances in understanding binding regions and critical contacts such as hotspots (Goncearenco et al. 2017; Ozdemir et al. 2019; Fuller, Burgoyne, and Jackson 2009). Therefore, investigating the properties of these interactions is critical for the identification of drug targets and alternative pathways for existing drugs (Lu et al. 2020). Structural clustering of interfaces is a strategy to create clusters with similar topology and limit the search space for possible candidates. Consequently, interfaces that belong to the same structural cluster may be filtered for repurposing a drug that already targets another protein in the same cluster. Further physicochemical analysis of the interfaces and binding sites helps to identify proteins that have the potential to be off-targets or new targets to the drug molecule can be identified.

TIMBAL (Higueruelo, Jubb, and Blundell 2013), 2P2Idb (Basse et al. 2016), iPPI-DB (Labbé et al. 2016) and DLIP (Ikeda et al. 2022) are a few public databases that have been made available thus far that contain chemical and target information linked to PPIs. TIMBAL is a resource for the automatic selection of small molecules modulating manually selected protein-protein complexes, whereas 2P2Idb is a hand-curated database of the structures of protein–protein complexes with known orthosteric inhibitors. iPPIdb also retrieves data manually from the literature. As of now, both TIMBAL and 2P2Idb are not accessible to the users. DLiP, on the other hand, is a database for facilitating the search for chemical structures and molecular descriptors related to PPI-oriented chemical libraries. It allows the search for newly-synthesized compounds designed to inhibit PPIs. It contains compound library data for PPI targets and known PPI inhibitor data collected from public databases. However, these databases have a small number of PPI targets, and the need for developing new tools is crucial.

Here, we present a curated large dataset for drug-like molecules in protein interfaces together with DiPPI (Drugs in Protein-Protein Interfaces), a publicly available website where users can access protein interfaces extracted from PDB and their associated drug-like molecules and FDA-approved drugs. Additionally, several physicochemical properties for the interfaces and drugs are presented on the webpage to guide users on the molecule and target selection. We have further investigated the relevance of our results using a docking procedure to find alternative drug-target pairs in a case study. Our work presents extensive protein interface data by analyzing 534,203 computationally derived interfaces and up-to-date drug-like molecules that are associated with these interfaces. DiPPI is freely available for data access and downloads at http://interactome.ku.edu.tr:8501. Users can download their query results and batch data files for interface and drug-like molecule associations.

## 2. Methods

DiPPI has two sites for protein interfaces and drug-like molecules (**Figure 1**). The datasets of protein interfaces and drug-like molecules are merged to identify drug-like molecules specifically targeting interfaces.

**Figure 1.**
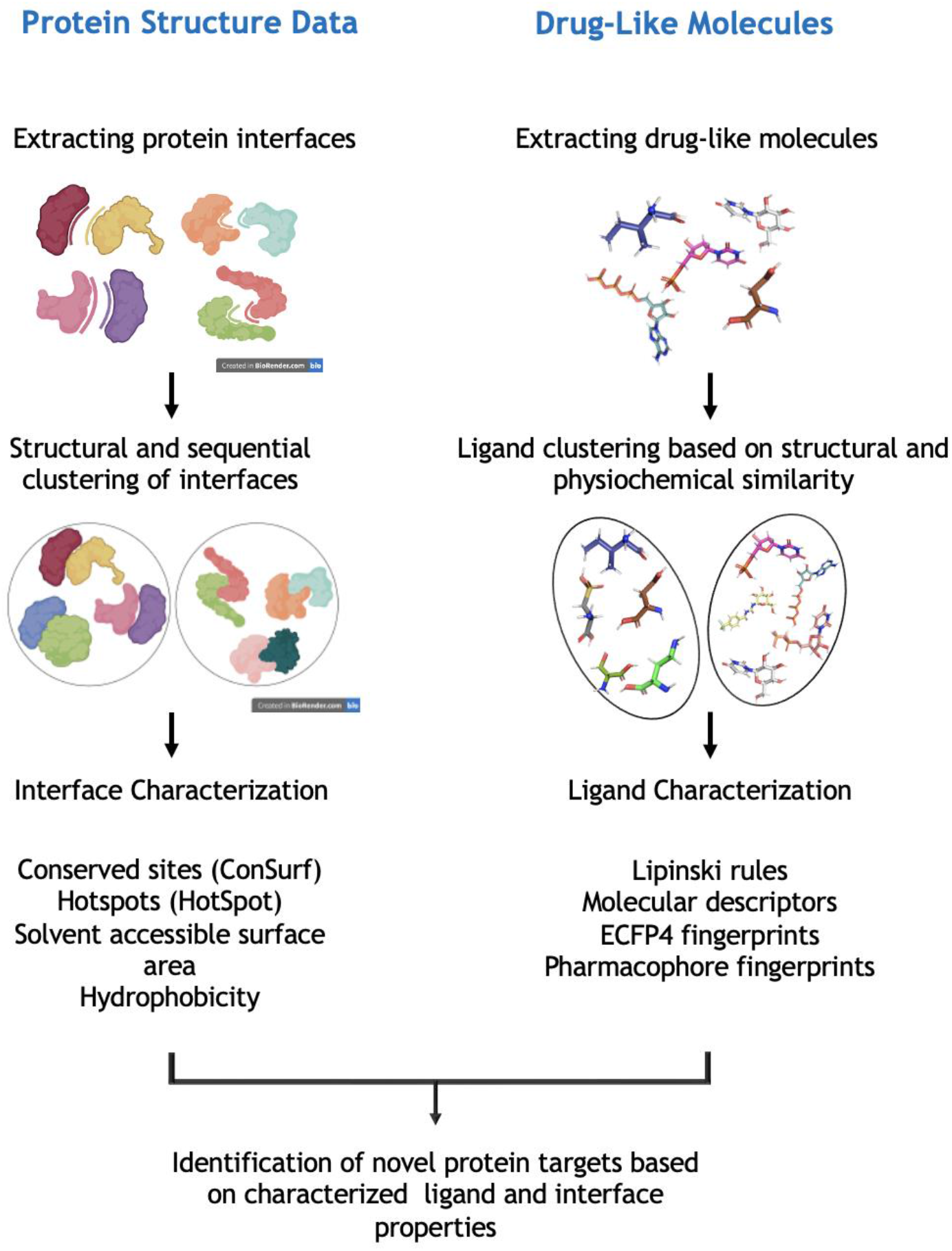
Flowchart of DiPPI process. Interface clusters are created as described in Abali et al., 2022. Drug-like small molecule data is curated from five databases. Relevant physicochemical properties are calculated for interface and drug-like molecules.

### 2.1. Creating interface clusters

Interface clusters are created as described in Abali 2021 (Abali 2021). The total set contains 534,203 interfaces from all available 3D structures in PDB as of March 2022. Interfaces are defined by two residues with atoms at most a certain distance (defined as vdW radii of involving atoms plus a threshold of 0.5 Å) away from each other. Interfaces are clustered by structural similarity using agglomerative hierarchical clustering. Representatives are selected as interfaces that are the most similar to all other interfaces in the same cluster.

### 2.2. Identification and Characterization of Small Drug-like Molecules

One of the critical steps in drug repositioning studies is the selection of small molecules or drug molecules to be repositioned. In this study, seven different databases, namely LigandExpo (Feng et al. 2004), ChEMBL (Gaulton et al. 2017), iPPIdb (Gaulton et al. 2017; Labbé et al. 2016), PDID (C. Wang et al. 2016), Open Targets Platform (Ochoa et al. 2021), BindingDB (Gilson et al. 2016) and ZINC database (Sterling and Irwin 2015), were used in the drug-like molecule data acquisition process. This dataset is further curated to eliminate cofactor, coenzyme, and ion molecules as annotated in PDBe (Mir et al. 2018).

### 2.3. Identification of drug-like molecules in the interfaces

After compiling the drug-like molecule data, we further selected drug-like small molecules associated with the proteins in the interface set. The interface dataset has 534,203 interfaces representing different chains of 98,632 PDB structures. 335,648 interfaces were found to belong to proteins associated with drug-like small molecules at any position; therefore, the downstream analyses were conducted using these interfaces. Although these proteins are characterized as binding to drug-like molecules, it is unclear if the bound drugs are located at the interfaces or any other protein region. As we aim to identify drug-like structures at the interfaces within the scope of this study, it is necessary to determine the exact positions where these molecules bind to the protein structure. To assess this, we obtained the binding positions of the drug-like molecules to the associated proteins through PDBsum (Laskowski et al. 1997) (http://www.ebi.ac.uk/pdbsum) database and mapped these binding positions to the interface residues. The following characterization steps used small molecules obtained at the end of this mapping process.

### 2.4. Dataset Characterization

For the characterization of drug-like molecules and FDA-approved drugs, various molecular descriptors are calculated with the Python RDKit module (2021_03_5), including but not limited to lipophilicity (logP), molecular weight, polar surface area (PSA), the number of rings, the number of carbons, the number of heteroatoms, the number of rotatable bonds (ROTB), the number of hydrogen bond acceptors (HBA), number of hydrogen bond donors (HBD), octanol-water partition coefficient (ALOGP), BalabanJ value and BertzCT indices. We further calculated whether these molecules are in agreement with the drug-likeliness criteria of Lipinski (Lipinski et al. 2001), Ghose (Ghose, Viswanadhan, and Wendoloski 1999), Veber (Ghose, Viswanadhan, and Wendoloski 1999; Veber et al. 2002), Egan (Egan, Merz, and Baldwin 2000), Muegge (Muegge 2002) as well as their quantitative estimate of drug-likeliness (QED) (Bickerton et al. 2012). Details of the selected criteria are given in Supplementary Files.

For the characterization of interfaces, the following properties are either calculated or collected from various databases: hotspots from HotPoint (Tuncbag, Keskin, and Gursoy 2010), conservation scores from ConSurf (Ben Chorin et al. 2020), basic characteristics (i.e., hydrophilicity, hydrophobicity, and polarity) of interface residues, and ligand binding energies from BindingDB (Gilson et al. 2016).

### 2.5. Small Molecule Clustering

In addition to clustering interfaces, we also clustered small molecules to exploit their similarities in the search for an alternative drug molecule. Structurally and physicochemically similar molecules are assumed to carry similar functions and be involved in similar processes (Bajorath 2017; Eckert and Bajorath 2007). We clustered drug-like small molecules according to their structural and pharmacological properties using molecular fingerprint analysis to limit the space in repurposing studies. In this direction, we extracted extended connectivity fingerprints (ECFP4) and pharmacophore fingerprints of small molecules using SMILES profiles and calculated fingerprint similarity with Tanimoto distance (Rogers and Hahn 2010; McGregor and Muskal 1999)(Bajusz, Rácz, and Héberger 2015; Rifaioglu et al. 2019)(Rogers and Hahn 2010; McGregor and Muskal 1999). ECFPs are known as effective and widely used fingerprints in drug discovery research and produce either better or comparable results to other fingerprint types in many applications (Bender et al. 2009; O’Boyle and Sayle 2016; Stepišnik et al. 2021; Menke and Koch 2021). Pharmacophore fingerprints can serve as an alternative to ECFP as they show the structural and chemical properties associated with the binding of molecules to other molecules.

After extracting fingerprints and quantifying their similarity, they are clustered using the Butina algorithm (Butina 1999), a preferred clustering method in the pharmacological context (Kutlushina et al. 2018; S. Wang et al. 2022; Zhou et al. 2022). When the results were evaluated regarding the distance within the clusters and the number of elements in the formed clusters, the appropriate threshold value was determined as 0.6 for ECFP4 and 0.5 for pharmacophore fingerprints, respectively **(Table S1**). A higher distance threshold means the molecules will be clustered into fewer clusters as more molecules will be similar. As the threshold decreases, clusters become smaller, eventually forming singletons.

Drug-like molecules appearing in the same cluster are provided on the website for each ligand to provide a guide to the users for prioritizing alternative drug-like molecules.

### 2.6. Docking For Case Study

In this study, we performed docking using AutoDock Vina (Eberhardt et al. 2021), and MGLTools (Morris et al. 2009) is used for the receptor and ligand preparation. The 3-dimensional structures of drugs at reference pH are downloaded from the ZINC database (Sterling and Irwin 2015). Additionally, BioPython (Cock et al. 2009) and NumPy (Harris et al. 2020) packages in Python and Open Babel (O’Boyle et al. 2011) are used. Protein-disease relation is found from DisGeNET (Piñero et al. 2017).

## 3. Results

### 3.1 Properties of Drug-Like Molecules

Filtered data from selected databases contains 11,011 small molecules known to have potential drug-like properties (see Section 2.2) and possess structure information in PDB (**Figure 2**). A list of selected drug-like small molecules and eliminated molecules can be found on the website.

**Figure 2.**
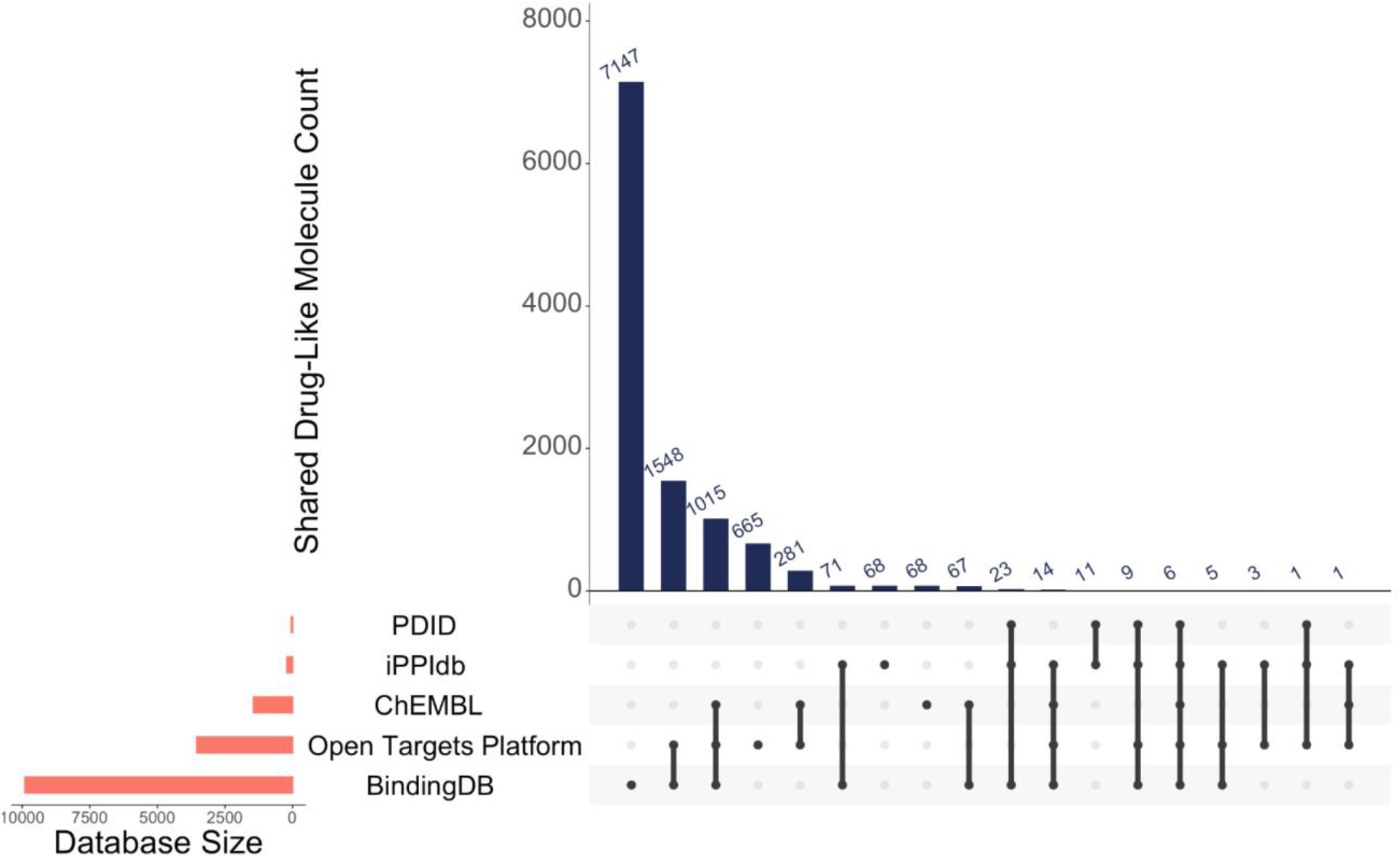
Distribution of drug-like small molecules with respect to source database

We also retrieved FDA-approved drugs from the ZINC database (Sterling and Irwin 2015). 335 out of 1,615 FDA-approved drugs are found to bind to an interface in our dataset. The list of these drugs can be found in the DiPPI GitHub repository. It can also be accessed through the DiPPI website.

### 3.1. Properties of Interfaces

In order to curate a dataset composed of the protein-protein interfaces and their bound drugs, we used an interface dataset extracted from all available structures in PDB (Abali et al. 2021) and drug-like small molecules from five databases. 2,214 out of 11,011 drug-like small molecules are found to be bound to at least one interface in our dataset of 335,648 interfaces. Interfaces are further filtered based on the FDA approval of the bound drug. As a result, 19,960 interfaces were found to have at least one FDA-approved drug associated with them. The resultant dataset is described in **Table S2**.

19,960 FDA-approved drug-bound interfaces cover a variety of proteins. 11,113 unique proteins corresponding to 2,378 Pfam families and 423 KEGG pathways are represented. The details for the ten most occurring KEGG pathways and Pfam families are given in **Table S3** and **Table S4**, respectively. Not surprisingly, overview pathways such as metabolic pathways (KEGG id: 01100), which are a combination of multiple pathways, are the most populated in our dataset (**Table S3**). In addition, some disease-related pathways are also highly populated, such as Alzheimer’s disease pathway (KEGG id: 05010) and the non-alcoholic fatty liver disease pathway (KEGG id: 04932). Some of the most populated Pfam families in our dataset are Immunoglobulin C1-set domain (Pfam id: PF07654) and Glycosyl hydrolases family 2 (Pfam id: PF00703) **Table S4**.

### 3.2. Ligand Clustering

Ligands are clustered into groups using ECFP4 and pharmacophore fingerprints. Members from an example cluster of ECFP4 and pharmacophore fingerprints can be seen in **Figure 3**. Molecules in the same cluster from both fingerprint analyses are given in the data table provided on the website. Users can process their results for either method upon download. As ECFP4 and pharmacophore fingerprints encode different characteristics regarding the ligand structure, using ECFP4 and pharmacophore fingerprints together will provide a better insight into their characteristics.

**Figure 3.**
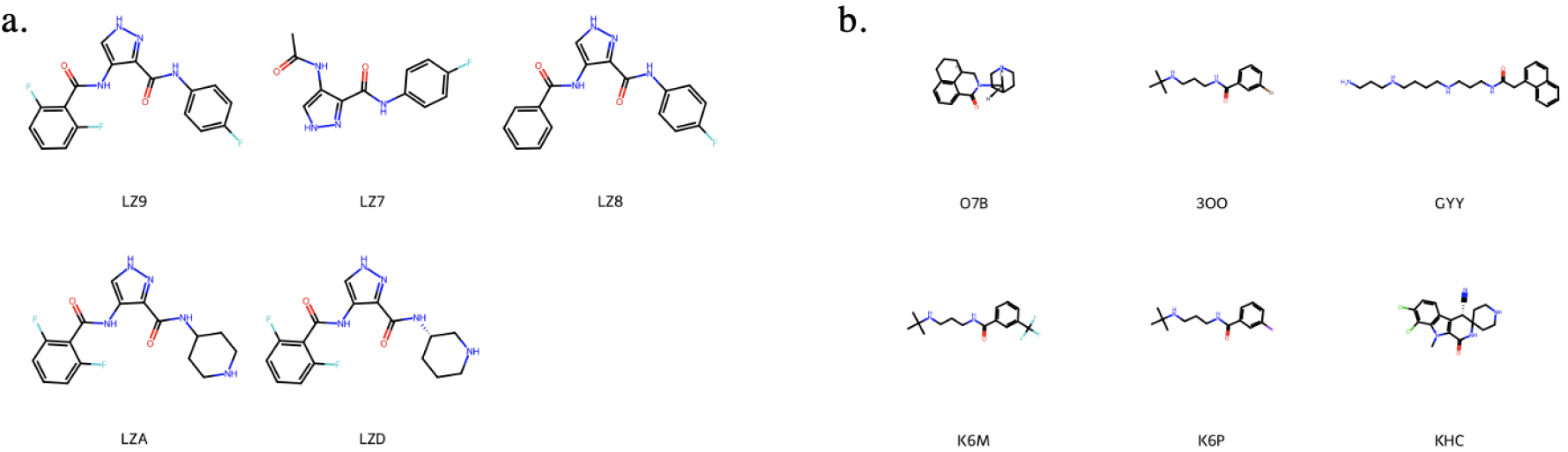
Members of a randomly selected cluster for (a) ECFP4 fingerprints (b) pharmacophore fingerprints.

### 3.3. Properties of Drugs Bound to Interfaces

Molecular descriptors of all drug-like molecules at interfaces are calculated and filtered based on the criteria from Lipinski, Ghose, Veber, Egan, Muegge, and QED. Of 11,011 interface-bound drug-like molecules, 7,905 of them have no violations of Lipinski’s rules, 4,799 of them have no violations of Ghose criteria, 9,595 of them have no violations of Veber criteria, 9,011 of them have no violations of Egan criteria, and 4,650 of them has no violations to Muegge criteria. Of 335 FDA-approved drugs which are bound to at least one interface, 228 of them have no violations of Lipinski’s rules, 103 of them has no violations of Ghose criteria, 275 of them have no violations of Veber criteria, 253 of them have no violations to Egan criteria and 123 of them has no violations to Muegge criteria. The distribution of some of the molecular descriptors and the distribution of QED scores of these drugs are given in **Figure S1**.

### 3.4. Website Overview

The DiPPI website has two modules for the users to submit their queries. In the ‘Query by Interface’ module, users can submit the PDB ID of the interface of interest, together with an optional selection of the desired ligand to generate a downloadable table (**Figure 4**). The table contains information regarding interface residues such as positions on the protein structure, position of the drug-like molecule on the interface, associated ligand ID, and hotpot residues if the drug-like molecule binds to them. In the ‘Query by Drug’ module, users can search using the ligand ID and retrieve a table with summary information about the drug and its associated interfaces (**Figure 5**). Upon clicking, the table shows detailed information about the selected ligands, including but not limited to calculated molecular descriptors and ligand cluster information. Tables on the pages are downloadable for further processing by the users.

**Figure 4.**
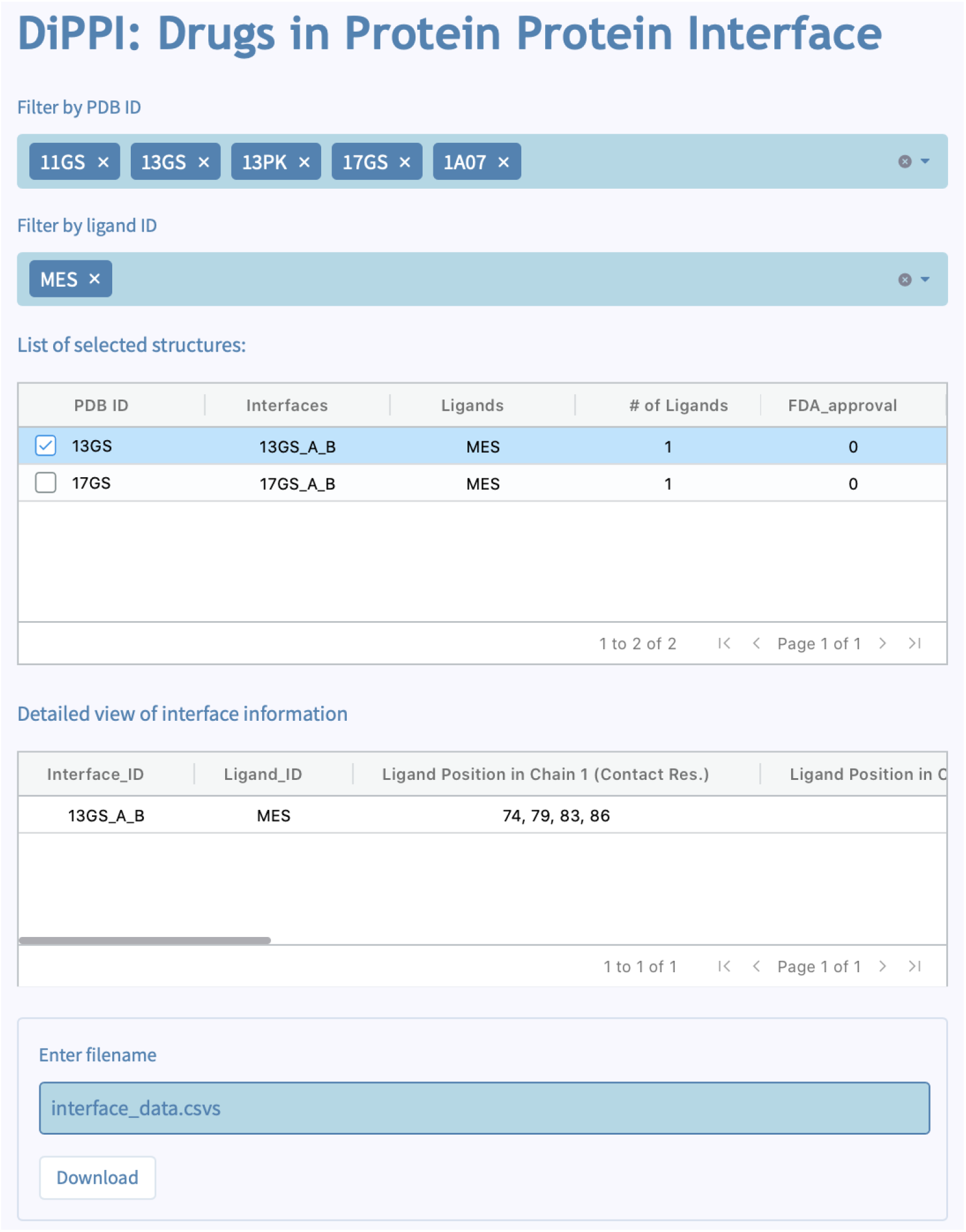
Example query for the ‘Query by Interface’ page in DiPPI.

**Figure 5.**
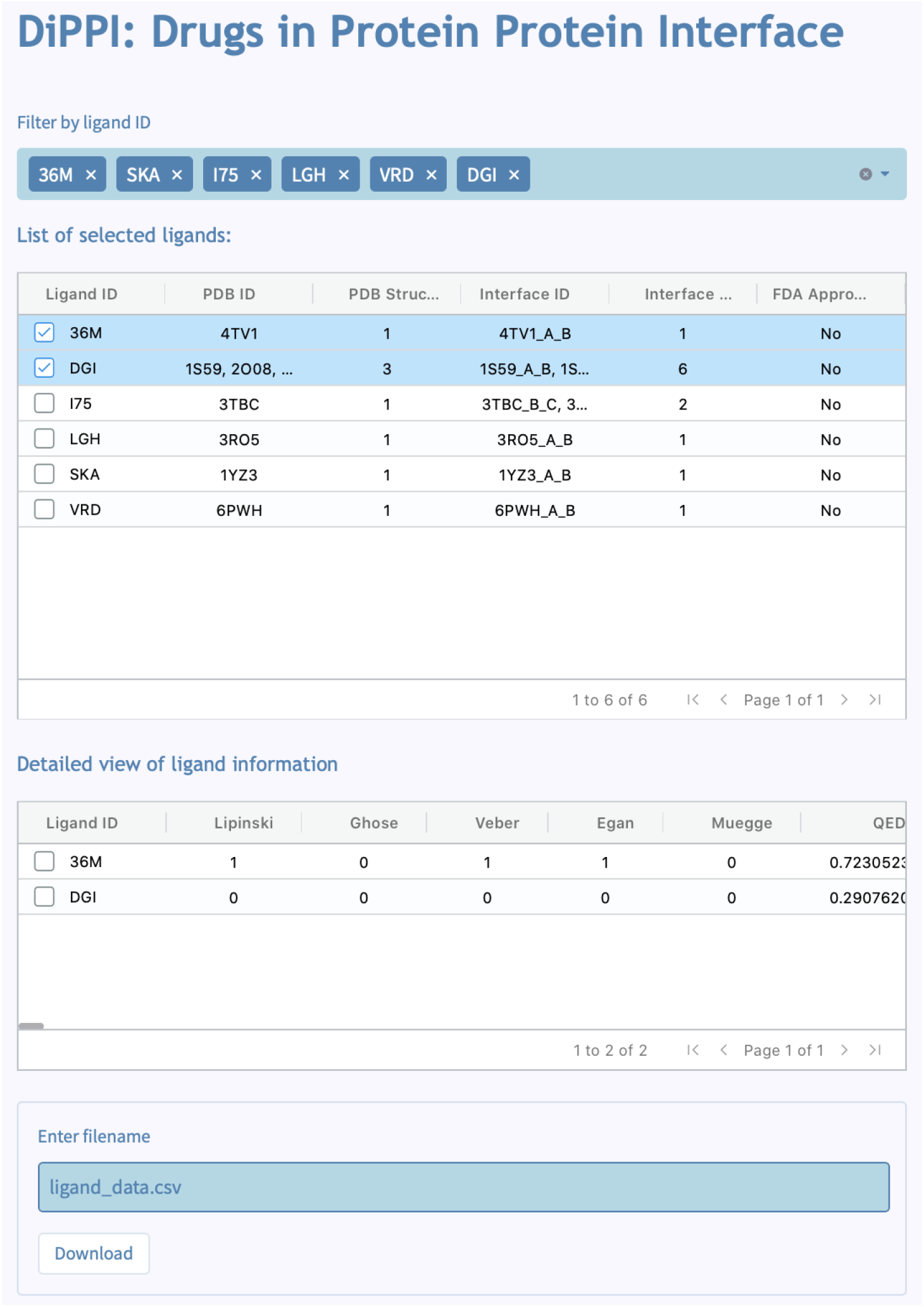
Example query for the ‘Query by Drug’ page in DiPPI.

### 3.5. Case Study

We tested the usability of our website by performing an *in-silico* novel drug identification for a protein interface which can further be validated *in vitro*. For this, we selected a set of proteins that have similar interfaces. In this set, there are 85 heterodimers, 12 of which have a drug in their interfaces. We docked the drugs to other proteins in the set. This selection is based on the number of unique FDA-approved drugs that bind to the interfaces.

The unique drugs in this cluster, their usage, and some of their molecular descriptors are given in **Table S5**. These drugs have a variety of uses, such as hypertension treatment and hypertriglyceridemia treatment. The proteins in the selected structural cluster and the diseases they are related to are given in **Table S6**. Protein-disease relation is found with DisGeNET (Piñero et al. 2017) The proteins within the structural cluster are found to be related to numerous diseases, including breast carcinoma, hypertensive disease, and liver-related diseases.

The drugs at the protein-protein interface are docked to other protein-protein interfaces in the same cluster, hence, to the structurally similar protein-protein interfaces. In total, there were 364 such cases, and results with favorable binding energy to at least one side of the protein-protein interface, i.e., lower than -7.75 kcal/mol (Nguyen et al. 2020), might be candidates for drug repurposing. Among these results, we will discuss the case where mifepristone binds to the interface between retinoid X receptor alpha (RXRA) and Nuclear Receptor Coactivator 2 (**Figure 6**). Mifepristone is a progesterone receptor antagonist used to terminate pregnancy up to 63 days gestation (Dunn and Brooks 2018). In our results, mifepristone binds to RXRA with an energy of -8.233 kcal/mol. RXRA is a member of the nuclear receptor superfamily (X. Ma et al. 2018). Type 2 nuclear receptors are found in the nucleus and regulate biological functions such as growth, differentiation, and metabolism (de Almeida and Conda-Sheridan 2019; R. M. Evans 1988). Nuclear receptor RXRA plays a role in transcriptional regulation (Ronald M. Evans and Mangelsdorf 2014) and has been considered as a target for cancer and metabolic diseases such as obesity and diabetes (Altucci et al. 2007).

**Figure 6.**
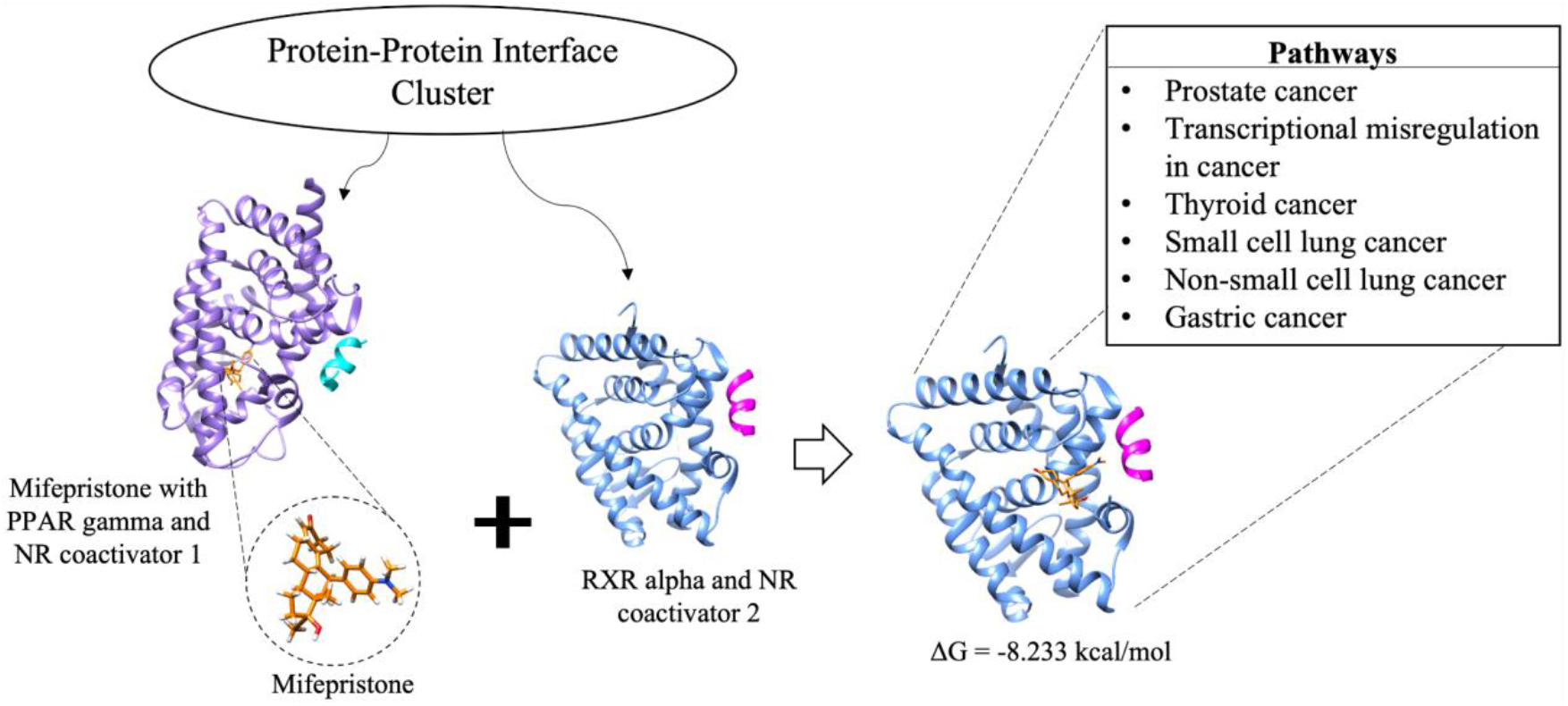
Mifepristone (orange) bound to the PPAR gamma-Nuclear receptor coactivator 1 interface is docked to the RXR alpha-Nuclear receptor coactivator 2 interface in the same protein-protein interface cluster found in DiPPI. As a result, mifepristone used for medical abortion is suggested to be repurposed for cancer treatment.

RXRA is involved in pathways such as thyroid cancer, PI3K-Akt pathway, Hepatitis C, pathways in cancer, small cell lung cancer, non-small cell lung cancer, and gastric cancer in KEGG (Kanehisa et al. 2017). In addition, RXRA is associated with prostatic neoplasms, according to DisGeNET. There are several studies for repurposing mifepristone as a cancer drug. A phase II study of mifepristone in castration-resistant prostate cancer (CRPC) suggested that mifepristone, when combined with an agent that prevents the increase in adrenal androgens, might benefit CRPC patients (Taplin et al. 2008; Turanli et al. 2018). Furthermore, another study reported the beneficial effects of mifepristone on murine testicular and prostate cancer (Check et al. 2010). In addition, mifepristone is suggested to be inducing apoptosis for androgen-independent prostate cancer cells (El Etreby, Liang, and Lewis 2000). Moreover, a combination of mifepristone and progesterone had an inhibitory effect on the growth of ovarian mesenchymal stem/stromal cells of females with *BRCA*^*1*−*/2*−^ mutation having a higher risk of ovarian cancer. Hence, it is proposed for ovarian cancer prevention (Ponandai-Srinivasan et al. 2019). In another study, inhibition of growth is observed in cancer cell lines from the nervous system, breast, prostate, ovary, and bone tissues with mifepristone by reducing the activity of Cdk2 (Tieszen et al. 2011). The binding of mifepristone to RXRA might be another mechanism that reduces the growth of cancer cells. Additionally, it is reported that highly metastatic cancer cells exhibit decreased migration and invasion when exposed to mifepristone (Ritch et al. 2019).

Although the proposed mechanism might differ, there is various supporting evidence in the literature for repositioning mifepristone in cancer treatment that might be explained by protein-protein interface structural similarities on the molecular level. This website provides the data for drug repurposing studies through protein-protein interface and drug clusters.

## 4. Conclusions

In this study, we present DiPPI, a website for the investigation of drug-like molecules in protein interfaces. Our website provides users with a curated collection of drug-like molecules and their computed properties, along with the characterization of interfaces that contain a drug-like molecule in downloadable format. In total, we investigated 534,203 predicted protein interfaces and 11,011 drug-like small molecules. Of all investigated molecules and interfaces, 335,648 interfaces are found to contain 2,214 drug-like molecules in total. Our website offers a guide to researchers who want to search for alternative targets and drugs in their drug repurposing studies by limiting the search space by allowing them to filter similar targets and drugs to their queries.

## 5. Acknowledgement

This project has been partially funded by TUSEB 4448/4081.

## Notes

### Competing Interest Statement

The authors have declared no competing interest.

