## Supplementary Material for "DiPPI: A curated dataset for drug-like molecules in protein-protein interfaces"

#### Supporting Information on Dataset Characterization

The molecular descriptors are filtered based on the following criteria:

1) Lipinski's Rule of Five (Lipinski et al. 2012):
   a) Lipophilicity (logP) < 5
   b) Molecular weight (MW) < 500
   c) The number of hydrogen bond donors (HBD) < 5
   d) The number of hydrogen bond acceptors (HBA) < 10
2) Ghose's criteria (Ghose, Viswanadhan, and Wendoloski 1999):
   a) 160 ≤ Molecular weight (MW) ≤ 480
   b) -0.4 ≤ Lipophilicity (logP) ≤ 5.6
   c) 40 ≤ Molar refractivity (MR) ≤ 130
   d) 20 ≤ The number of atoms ≤ 70
3) Veber's criteria (Veber et al. 2002):
   a) The number of rotatable bonds ≤ 10
   b) Polar surface area (PSA) ≤ 140
4) Egan's criteria (Egan, Merz, and Baldwin 2000) (ref):
   a) Lipophilicity (logP) ≤ 5,88
   b) Polar surface area (PSA) ≤ 131,6
5) Muegge's criteria (Muegge 2002):
   a) 200 ≤ Molecular weight (MW) ≤ 600
   b) -2 ≤ Lipophilicity (logP) ≤ 5
   c) Polar surface area (PSA) ≤ 150
   d) The number of rings ≤ 7
   e) The number of carbons > 4
   f) The number of heteroatoms > 1
   g) The number of rotatable bonds (ROTB) ≤ 15
   h) The number of hydrogen bond acceptors (HBA) ≤ 10
   i) The number of hydrogen bond donors (HBD) ≤ 5
6) Quantitative estimate of drug-likeness (QED) (Bickerton et al. 2012):
   a) QED score is calculated with Python's RDKit module. QED score uses the following parameters: octanol-water partition coefficient (ALOGP), molecular weight (MW), the number of hydrogen bond acceptors (HBA), the number of hydrogen bond donors (HBD), polar surface area (PSA), rotatable bond count (ROTB), aromatic ring count (AROM), the presence of unwanted chemical functionalities / structural alerts (ALERTS)

### Figures

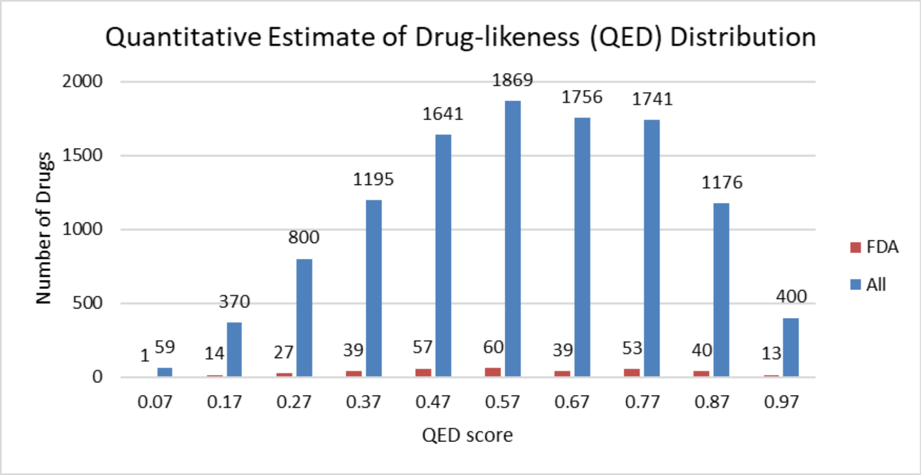
Quantitative Estimate of Drug-likeness (QED) Distribution
Number of Drugs
0
500
1000
1500
2000
1
59
14
370
27
800
39
1195
57
1641
60
1869
39
1756
53
1741
40
1176
13
400
0.07
0.17
0.27
0.37
0.47
0.57
0.67
0.77
0.87
0.97
QED score
FDA
All

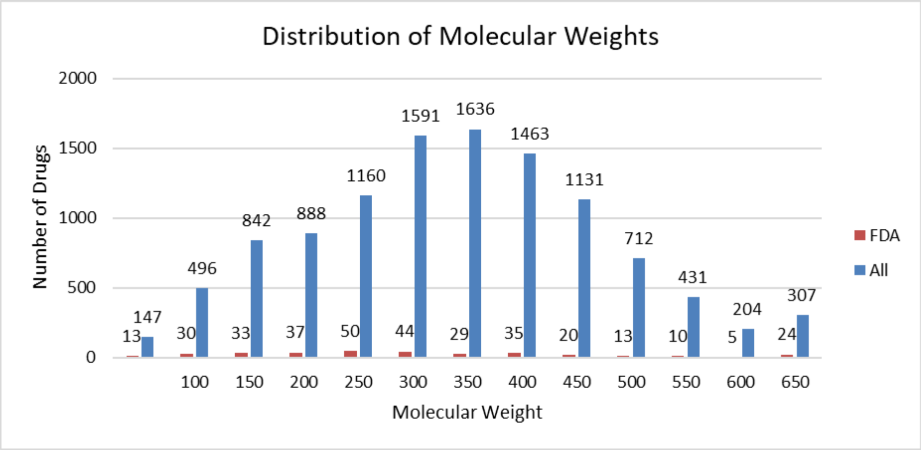
Distribution of Molecular Weights
Number of Drugs
0
500
1000
1500
2000
13
147
30
496
33
842
37
888
50
1160
44
1591
29
1636
35
1463
20
1131
13
712
10
431
5
204
24
307
100
150
200
250
300
350
400
450
500
550
600
650
Molecular Weight
FDA
All

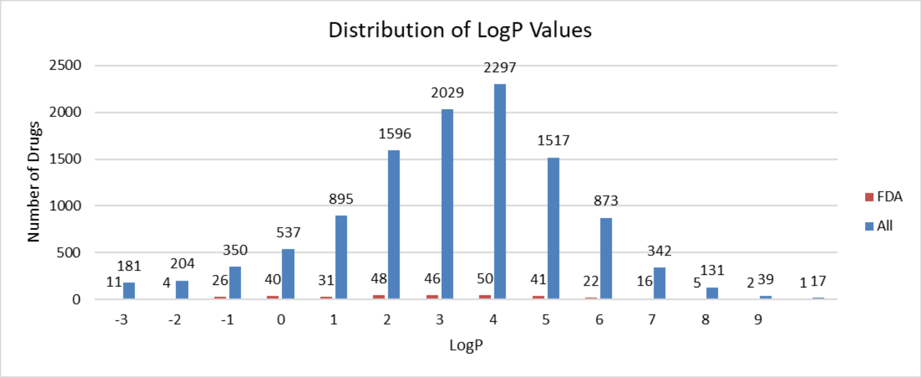
Distribution of LogP Values
Number of Drugs
0
500
1000
1500
2000
2500
11
181
4
204
26
350
40
537
31
895
48
1596
46
2029
50
2297
41
1517
22
873
16
342
5
131
2
39
1
17
-3
-2
-1
0
1
2
3
4
5
6
7
8
9
LogP
FDA
All

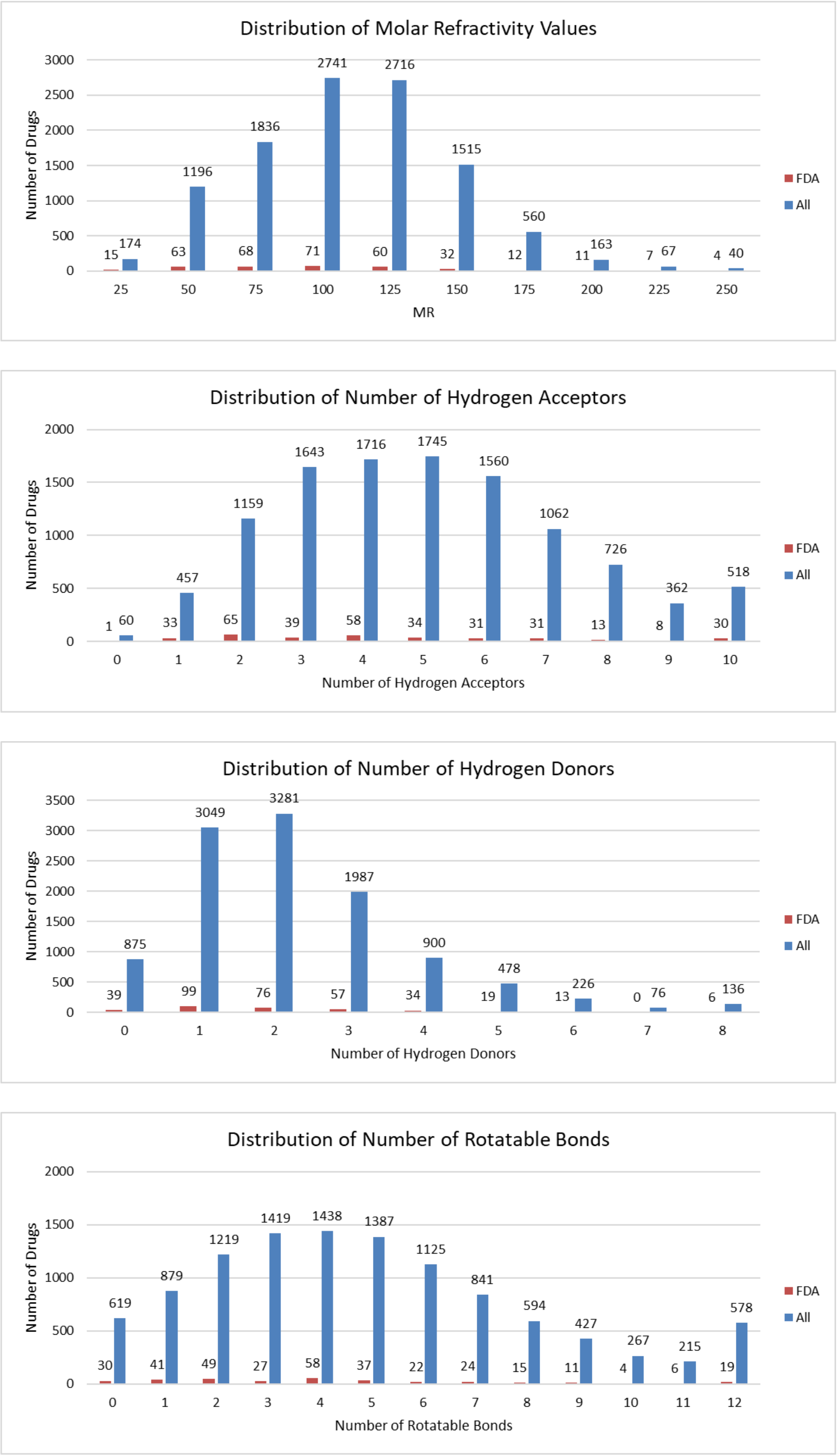

**Figure S1.** Distribution of Lipinski's rules related to molecular descriptors of FDA-approved drugs.

#### Tables

**Table S1.** Ligand cluster characterization for ECFP4 and pharmacophore fingerprints

| | ECFP4 | Pharmacophore |
|---|---|---|
| Selected threshold value | 0.6 | 0.5 |
| Total cluster number | 6084 | 2799 |
| Clusters with only one molecule | 4270 | 1473 |
| Clusters with more than 5 molecules | 242 | 383 |
| Clusters with more than 10 molecules | 53 | 162 |
| Clusters with more than 25 molecules | 10 | 51 |
| Clusters with more than 50 molecules | 1 | 23 |
| Molecule count in the most crowded cluster | 93 | 303 |
| Similarity between two random molecules within a cluster | 0.66 | 0.65 |
| Similarity between two random molecules in two random clusters | 0.07 | 0.16 |

**Table S2.** The description of the dataset

| | |
|---|---|
| Number of investigated proteins | 98,632 |
| Number of investigated interfaces | 534,203 |
| Number of interfaces belonging to proteins with bound drugs (any region) | 335,648 |
| Number of interfaces to which at least one drug-like molecule binds to | 53,452 |
| Number of interfaces to which at least one FDA-drug binds to | 19,960 |
| Number of investigated drug-like small molecules | 11,011 |
| Number of eliminated small molecules | 402 |
| Number of investigated FDA-approved drugs | 1,615 |
| Number of drug-like small molecules that bind to at least one interface residue | 2,214 |
| Number of FDA-approved drugs that bind to at least one interface residue | 335 |

**Table S3.** Top 10 most occurring KEGG pathways in FDA-approved drug-bound interfaces dataset

| **KEGG Pathway Name** | **KEGG Pathway ID** | **Number of Occurences** |
|---|---|---|
| Metabolic pathways | 01100 | 9342 |
| Biosynthesis of secondary metabolites | 01110 | 2524 |
| Alzheimer disease | 05010 | 1888 |
| Biosynthesis of cofactors | 01240 | 1770 |
| Microbial metabolism in diverse environments | 01120 | 1543 |
| Diabetic cardiomyopathy | 05415 | 1456 |
| Pathways of neurodegeneration - multiple diseases | 05022 | 1446 |
| Non-alcoholic fatty liver disease | 04932 | 1324 |
| Type I diabetes mellitus | 04940 | 1310 |
| Human T-cell leukemia virus 1 infection | 05166 | 1275 |

**Table S4.** Top 10 most occurring Pfam families in FDA-approved drug-bound interfaces dataset

| **Pfam Family Name** | **Pfam ID** | **Number of Occurences** |
|---|---|---|
| Immunoglobulin C1-set domain | PF07654 | 1794 |
| Reverse transcriptase connection domain | PF06815 | 924 |
| Reverse transcriptase thumb domain | PF06817 | 924 |
| Reverse transcriptase (RNA-dependent DNA polymerase) | PF00078 | 924 |
| Glycosyl hydrolases family 2, TIM barrel domain | PF02836 | 818 |
| Glycosyl hydrolases family 2 | PF00703 | 816 |
| Glycosyl hydrolases family 2, sugar binding domain | PF02837 | 816 |
| Beta galactosidase small chain | PF02929 | 720 |
| Beta-galactosidase, domain 4 | PF16353 | 720 |
| Neurotransmitter-gated ion-channel ligand binding domain | PF02931 | 660 |

**Table S5.** The drugs docked in the case study, their usage, and the following molecular descriptors: molecular weights (MWt), the number of hydrogen acceptors (HA), the number of hydrogen donors, and the number of rotatable bonds.

| Drug Name | Ligand ID | Usage | MWt | logP | HA | HD | ROTB |
|---|---|---|---|---|---|---|---|
| Bexarotene | 9RA | Used for the treatment of the skin manifestations of CTCL | 348.49 | 6.10 | 1 | 1 | 3 |
| Alitretinoin | 9CR | Used for the treatment of the lesions in patients with AIDS-related Kaposi's sarcoma | 300.44 | 5.60 | 1 | 1 | 5 |
| Mifepristone | 486 | Used to terminate intrauterine pregnancy | 429.60 | 5.41 | 3 | 1 | 2 |
| Docosahexaenoic acid | HXA | An omega-3 fatty acid | 328.50 | 6.55 | 1 | 1 | 14 |
| Fenofibric acid | F5A | Supplementary to the treatment of hypertriglyceridemia and high cholesterol | 318.75 | 3.81 | 3 | 1 | 5 |
| Telmisartan | TLS | Used for the treatment of hypertension | 514.63 | 7.26 | 5 | 1 | 7 |
| Eicosapentaenoic Acid | EPA | Related to Icosapent which is used in the treatment of hyperglyceridemia | 302.46 | 5.99 | 1 | 1 | 13 |

**Table S6.** The proteins that are in the structural cluster (with the cluster representative 5AZT_A_C) and the diseases they are the most related to. Only the diseases with DisGeNET scores higher than or equal to 0.5 are given.

| **Protein** | **Protein Name** | **Uniprot ID** | **Disease** | **Drug Bound Interfaces within the Cluster** |
|---|---|---|---|---|
| NCOA1 | Nuclear receptor coactivator 1 | Q15788 | Breast carcinoma | 3QT0_A_C, 1FM9_A_B, 1FM6_A_B, 1K74_A_B, 7BQ4_A_B, 7BQ0_A_B, 3VN2_A_C |
| NCOA2 | Nuclear receptor coactivator 2 | Q15596 | No results found with high scores | 3OAP_A_B, 4NQA_A_C, 4K6I_A_B, 1MV9_A_B |
| Nr0b2 | Nuclear receptor subfamily 0 group B member 2 | P97947 | No results found | This protein is in the cluster but none of the drugs are bound to its interfaces |
| RXRA | Retinoic acid receptor RXR-alpha | P19793, P28700 | Prostatic neoplasms | 1FM9_A_B, 1FM6_A_B, 1K74_A_B, 1XLS_A_I, 3OAP_A_B, 4NQA_A_C, 4K6I_A_B, 1MV9_A_B, 1XDK_A_C |
| RXRB | Retinoic acid receptor RXR-beta | P28702 | No results found with high scores | This protein is in the cluster but none of the drugs are bound to its interfaces |
| Med1 | Mediator of RNA polymerase II transcription subunit 1 | Q925J9 | No results found | 1XDK_A_C |
| PPARA | Peroxisome proliferator-activated receptor alpha | Q07869 | Fatty liver, liver neoplasms, hypertensive disease, malignant neoplasm of liver, reperfusion injury | 7BQ4_A_B, 7BQ0_A_B |
| PPARG | Peroxisome proliferator-activated receptor gamma | P37231 | Obesity, Familial Partial Lipodystrophy | 3QT0_A_C, 3VN2_A_C |

| | | | |
|---|---|---|---|
| | | | (Type 3),<br>Diabetes Mellitus (Non-Insulin-Dependent),<br>Hypertensive disease, Malignant tumor of colon,<br>Diabetic Nephropathy, Familial partial lipodystrophy,<br>Inflammation,<br>Diabetes Mellitus,<br>Atherosclerosis,<br>Colorectal Carcinoma,<br>Acute kidney injury,<br>Acute Lung Injury,<br>Glomerulonephritis,<br>Familial Partial Lipodystrophy Type 1),<br>Carotid Intimal Medial Thickness 1 |
